# Preservation of Chromatin Organization after Acute Loss of CTCF in Mouse Embryonic Stem Cells

**DOI:** 10.1101/118737

**Authors:** Naoki Kubo, Haruhiko Ishii, David Gorkin, Franz Meitinger, Xiong Xiong, Rongxin Fang, Tristin Liu, Zhen Ye, Bin Li, Jesse R. Dixon, Arshad Desai, Huimin Zhao, Bing Ren

**Affiliations:** Ludwig Institute for Cancer Research, La Jolla, CA 92093, USA; Department of Chemical and Biomolecular Engineering, University of Illinois at Urbana-Champaign, Urbana, IL 61801, USA; Salk Institute for Biological Studies, 10010 North Torrey Pines Road, La Jolla, CA 92037, USA; Department of Cellular and Molecular Medicine, University of California San Diego, La Jolla, CA 92093, USA; Department of Biochemistry, University of Illinois at Urbana-Champaign, Urbana, IL 61801, USA; Departments of Chemistry, Biochemistry, and Bioengineering, and Carl R. Woese Institute for Genomic Biology, University of Illinois at Urbana-Champaign, Urbana, IL 61801, USA; Department of Cellular and Molecular Medicine, Moores Cancer Center and Institute of Genome Medicine, UCSD School of Medicine, 9500 Gilman Drive, La Jolla, CA 92093, USA

## Abstract

The CCCTC-binding factor (CTCF) is widely regarded as a key player in chromosome organization in mammalian cells, yet direct assessment of the impact of loss of CTCF on genome architecture has been difficult due to its essential role in cell proliferation and early embryogenesis. Here, using auxin-inducible degron techniques to acutely deplete CTCF in mouse embryonic stem cells, we show that cell growth is severely slowed yet chromatin organization remains largely intact after loss of CTCF. Depletion of CTCF reduces interactions between chromatin loop anchors, diminishes occupancy of cohesin complex genome-wide, and slightly weakens topologically associating domain (TAD) structure, but the active and inactive chromatin compartments are maintained and the vast majority of TAD boundaries persist. Furthermore, transcriptional regulation and histone marks associated with enhancers are broadly unchanged upon CTCF depletion. Our results suggest CTCF-independent mechanisms in maintenance of chromatin organization.

## INTRODUCTION

Mammalian chromosomes reside in separate chromosome territories in interphase nuclei and are partitioned into topologically associating domains (TADs) characterized by strong intra-domain interactions and comparatively weak inter-domain contacts (Dixon et al., 2012; Nora et al., 2012). TADs are generally invariant across different cell types and evolutionarily conserved in related species (Dixon et al., 2012; Schmitt et al., 2016; Vietri Rudan et al., 2015). Cell-type specific chromatin interactions between distal *cis* regulatory elements and their target genes are largely constrained by the TADs. Some TADs are further divided into structures such as sub-TADs and insulated neighborhoods (Hnisz et al., 2016; Phillips-Cremins et al., 2013). TADs have been shown to allow enhancers to act on genes within the same domain (Symmons et al., 2016; Symmons et al., 2014), providing a mechanism for long-range control by distal regulatory elements (Dixon et al., 2016; Schwarzer and Spitz, 2014). Disruption of TADs and sub-domains has been shown to result in ectopic enhancer/promoter interactions and altered gene expression programs in cancers (Flavahan et al., 2016; Hnisz et al., 2016) and developmental disorders (Andrey et al., 2013; Franke et al., 2016; Lupianez et al., 2015). Understanding the molecular mechanisms that establish and maintain TADs and other features of chromatin organization will have significant implications in the study of a wide variety of human diseases.

The CCCTC-binding factor (CTCF) is widely believed to play a critical role in genome organization in bilaterian animals (Dekker and Mirny, 2016; Denker and de Laat, 2016; Ghirlando and Felsenfeld, 2016; Hnisz et al., 2016; Ong and Corces, 2014; Vietri Rudan and Hadjur, 2015). This ubiquitously expressed DNA binding protein is associated with a large number of genomic regions via a consensus sequence in the mammalian genome (Kim et al., 2007; Schmidt et al., 2012; Vietri Rudan and Hadjur, 2015; Xie et al., 2007). The CTCF binding sites are highly enriched at boundaries of TADs, sub-TADs, insulated neighborhoods, and at loop anchors (Dixon et al., 2012; Hnisz et al., 2016; Phillips-Cremins et al., 2013; Rao et al., 2014). Furthermore, the CTCF binding motifs at loop anchors and TAD boundaries preferentially adopt a convergent orientation, highlighting a role for the protein in the formation of TADs and chromatin loops (Guo et al., 2015; Rao et al., 2014). In support of this model, genomic deletions encompassing TAD boundaries which contain CTCF binding sites lead to a gain of interactions between neighboring TADs (Lupianez et al., 2015; Narendra et al., 2016; Narendra et al., 2015; Nora et al., 2012). Targeted disruption of specific CTCF binding motifs alters chromatin loops (de Wit et al., 2015; Guo et al., 2015; Narendra et al., 2016; Narendra et al., 2015; Sanborn et al., 2015). Moreover, knockdown of CTCF using RNA interference leads to an increase in contacts across TAD boundaries, and a reduction in contacts between putative CTCF-bound loop anchors (Zuin et al., 2014). Finally, comparison of chromatin domain organization in four mammalian species has revealed a close association between divergent CTCF binding sites and the gain or loss of TAD domains during evolution, further highlighting a conserved role of CTCF in genome organization (Vietri Rudan et al., 2015).

Despite the weight of evidence linking CTCF to TADs and chromatin loops, how CTCF shapes chromatin structure remains incompletely understood. The prevailing model postulates that CTCF acts together with the Cohesin complex, a ring-shaped structure consisting of Smc1, Smc3, Rad21 and SA1/2 subunits that mediate cohesion of sister chromatids during mitosis (Dekker and Mirny, 2016; Denker and de Laat, 2016; Ghirlando and Felsenfeld, 2016; Hnisz et al., 2016; Merkenschlager and Odom, 2013; Nativio et al., 2009; Ong and Corces, 2014; Vietri Rudan and Hadjur, 2015). This model is supported by the observations that the Cohesin complex nearly always co-localizes with CTCF in the genome (Hadjur et al., 2009; Parelho et al., 2008; Stedman et al., 2008; Wendt et al., 2008), that CTCF can physically associate with Cohesin (Xiao et al., 2011), that association of Cohesin with DNA depends on CTCF (Wendt et al., 2008), and that depletion of Cohesin leads to loss of long-range chromatin interactions (Nativio et al., 2009; Seitan et al., 2013; Sofueva et al., 2013; Zuin et al., 2014). The preferential convergent orientation of CTCF binding sites at loop anchors has led to the “extrusion” model in which a pair of tethered Cohesin complexes traversing the chromatin fiber in opposite directions pause at CTCF binding sites, leading to extrusion of the intervening DNA to form a chromatin loop and presumably chromatin domains (Fudenberg et al., 2016; Sanborn et al., 2015).

While the above model can explain the phenotypic consequences after disruption of CTCF binding motifs or inversion of CTCF binding sites (de Wit et al., 2015; Guo et al., 2015; Narendra et al., 2016; Narendra et al., 2015; Sanborn et al., 2015), many exceptions to the model have been reported. Notably, there is not a straightforward correspondence between the presence of CTCF binding sites and formation of a TAD boundary. In fact, roughly 15% of all TAD boundaries show no evidence of CTCF binding (Dixon et al., 2012). Further, nearly a third of chromatin loops anchored on CTCF binding sites show tandem CTCF motifs arrangement (Tang et al., 2015). In addition, although CTCF knockdown increases the frequency of contacts between neighboring TADs, overall TAD structure remains largely unchanged in these experiments (Zuin et al., 2014). The effects of CTCF knockdown become clear only after careful quantitative analysis. In these experiments RNA interference achieved about 80% depletion of CTCF protein levels, and the mild effect on chromatin organization was attributed to incomplete loss of CTCF protein after knockdown. Thus, a major question about the role of CTCF in chromatin organization remains unanswered: what happens to TADs and chromatin loops when cells are fully depleted of the CTCF protein?

Numerous attempts have been made to knock out the CTCF gene in mammalian cells, but the early embryonic lethality of CTCF null animals or growth inhibition in conditional knockout mouse ES cells have hindered further studies (Heath et al., 2008; Sleutels et al., 2012). In order to study chromatin organization after loss of CTCF, we employed the auxin-inducible degron (AID) system to acutely and nearly completely deplete the CTCF protein (Holland et al., 2012; Nishimura et al., 2009; Nishimura and Kanemaki, 2014). To our surprise, we found that the TAD structure is largely maintained after CTCF depletion. There is mild overall reduction of chromatin interaction frequencies, loss of Cohesin complex occupancy, and weakening of the strengths of TAD boundaries, but a large majority of TAD boundaries is preserved. The active and inactive chromatin compartments are also unaltered in the absence of CTCF, along with transcription profiles and histone modifications. Interestingly, TADs in the Lamina Associated Domains (LADs) exhibit a higher degree of weakening than in non-LAD regions. These results provide new insights into mechanisms that regulate chromatin architecture in mammalian cells.

## RESULTS

### Removal of CTCF binding on the genome by the auxin-inducible degron system

We genetically engineered a hybrid mouse embryonic stem cell (mESC) line F123 (129/CAST) (Gribnau et al., 2003) by inserting a sequence encoding the AID domain into 3’ end of the endogenous *Ctcf* gene using CRISPR/Cas9 and microhomology-mediated end-joining techniques (Nakade et al., 2014). The resulting cell clones were further engineered to stably express Tir1, which binds to CTCF-AID upon auxin exposure to induce rapid degradation of the CTCF-AID protein (Figure 1A: development of CTCF-AID mESCs). In this system, degradation of protein can be observed within few hours, and we confirmed the virtually complete depletion of CTCF within 24 hours of auxin exposure by Western blot (Figure 1B). The mESCs with AID-tagged CTCF can expand normally and auxin itself has no toxicity (Figure S1A). We observed that CTCF depleted cells could form colonies and were viable, but their growth was severely retarded (Figure 1C and Figure S1B). To ensure that the cells have enough time to go through several cell cycles without CTCF, we investigated the CTCF depleted cells for up to 4 days after auxin treatment. Additionally, to overcome potential clonal variations, we carried out in-depth molecular characterization using two independently derived cell clones.

We first performed CTCF ChIP-seq in each cell clone to confirm that CTCF occupancy is lost after auxin treatment at each time point (Figure 1D and 1E). The distribution of CTCF (p-value < 1 }10^−5^) in untreated CTCF-AID cells resembles that of normal ES cells (Shen et al., 2012), indicating that AID fusion does not affect its DNA binding (Figure S1C). The number of peaks dramatically decreased to less than 7% and 2 % of that in untreated CTCF-AID cells after 24 hours of auxin treatment in clone 1 and 2, respectively, and reached the minimum of less than 1% in both clones by 48 hours of auxin treatment (Figure 1F and Figure S1D-E). The remaining CTCF peaks in the CTCF depleted cells also exhibited significantly decreased enrichment levels. These results indicate that CTCF is nearly completely removed from the genome after 24 hours of auxin treatment and the penetrant depletion efficiency was maintained in both clones.

CTCF depleted cells grew significantly slower than untreated cells, but otherwise exhibited normal cycling behavior (Figure S1F), in contrast to previous observations of cell cycle arrest in T cells (Heath et al., 2008) and inability to proliferate of mouse ES cells after complete KO (Sleutels et al., 2012). Interestingly, the two CTCF-AID cell clones exhibit different growth rate both prior to and after CTCF depletion (Figure 1F). Nevertheless, judging from the Western blots and ChIP-seq analysis results, severe enough depletion of CTCF was achieved in both clones that allowed us to investigate the contribution of CTCF to chromatin architecture on a genome-wide scale.

**Figure 1.**
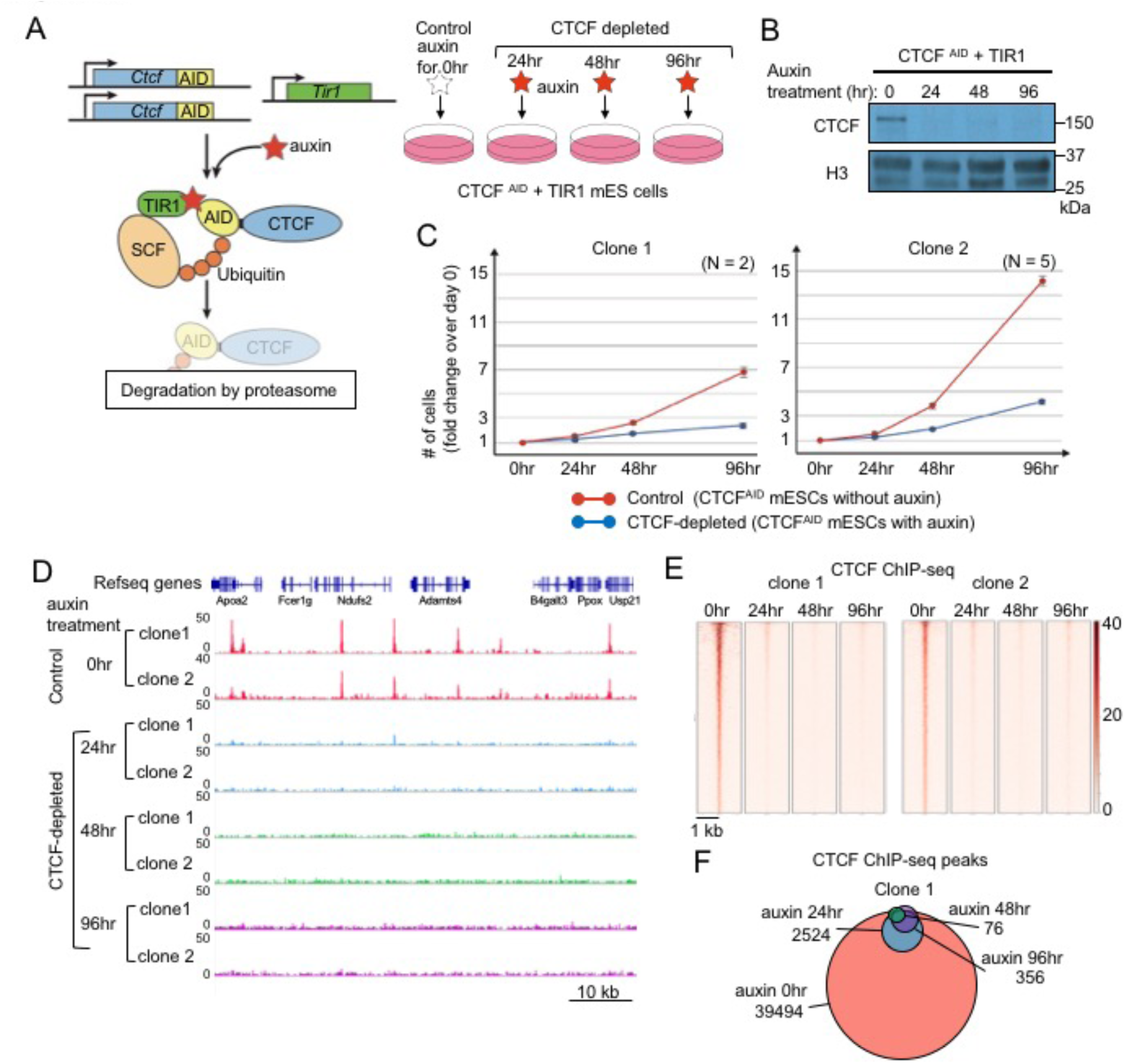
Acute depletion of CTCF in mESC by the auxin-inducible degron system. (A) Schematic representation of the generation of AID system in mESC and study design. (B) Western blot showing the depletion of CTCF protein after 24, 48, and 96 hours of auxin treatment. (C) Cell growth of control cells and CTCF-depleted cells treated with auxin for 4 days. (D) Snapshots of genome browser of two clones of CTCF ChIP-seq experiments showing that CTCF peaks observed in control cells were removed in CTCF-depleted cells. (E) CTCF ChIP-seq centered at all regions of CTCF peaks identified in the control (no auxin treatment) cells, and CTCF occupancy at the same regions in CTCF depleted cells at each time point. The CTCF peaks disappeared almost completely after auxin treatment in both two clones. (F) Venn-diagram comparing the numbers of CTCF ChIP-seq peaks identified in the control and CTCF-depleted cells in clone 1.

### TADs and compartments are largely maintained upon CTCF depletion

To examine the effect of CTCF depletion on chromatin organization, we performed *in situ* Hi-C experiments (Rao et al. 2014) with cells treated with auxin to trigger CTCF degradation for 0, 24, 48, and 96 hours. Hi-C reveals regions of genomic DNA that are in close spatial proximity in a genome-wide fashion using high-throughput sequencing. Surprisingly, TAD structure was maintained for several days after CTCF depletion and genome-wide loss of CTCF binding (Figure 2A). The vast majority of TAD boundaries persisted days after the near-complete loss of CTCF protein (Figure 2B), and the frequency of interaction within TADs (intra-TAD) as well as interaction across TADs (inter-TAD) (Krijger et al., 2016) was also preserved (Figure 2C). We did observe a slight weakening of TADs after CTCF depletion as evidenced by blurred edges in Hi-C heatmaps (Figure 2A), marginal decrease of insulation score (Figure 2D), increased interaction frequency across TAD boundaries (Figure S2A), and reduced Directionality Index (DI) adjacent to TAD boundaries (Figures 2E and S2B). However, despite this quantitative weakening of TADs, the degree to which TADs remain intact in the absence of CTCF is unexpected. These data demonstrate that CTCF is not strictly required for the maintenance of TADs.

**Figure 2.**
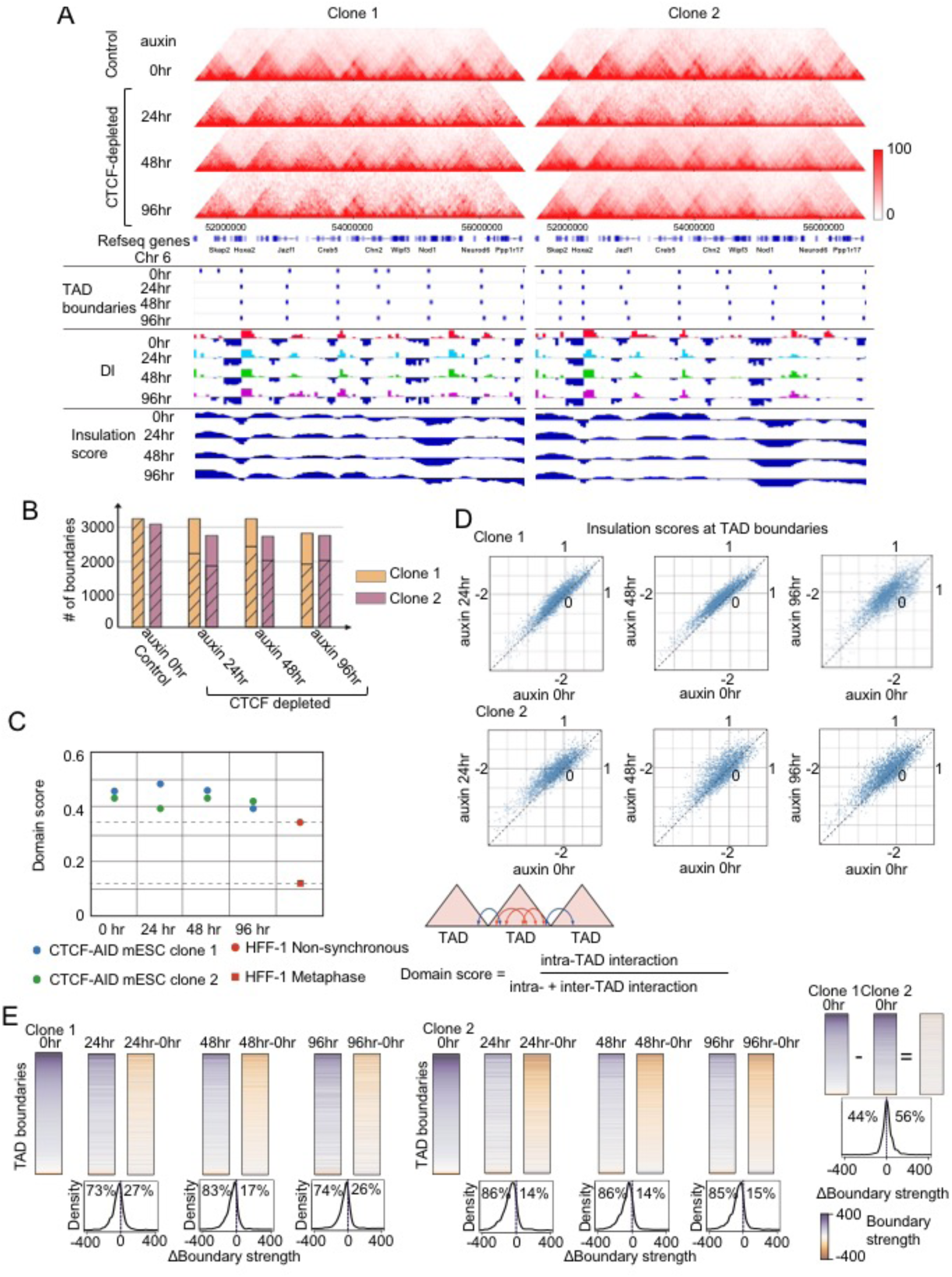
TAD structure and chromatin compartments are generally preserved after CTCF depletion. (A) Chromatin interaction heat maps in control and CTCF-depleted cells. The TAD boundaries, directionality index (DI), insulation score and CTCF ChIP-seq data in each clone are shown under the heatmap. (B) The number of TAD boundaries in control and CTCF depleted cells, and the hatched bars indicate the number of boundaries that overlapped with boundaries in control cells. (C) Frequency of interaction within TADs (intra-TAD) against interaction across TADs (inter-TAD) is shown as domain score (Krijger et al., 2016) in control and CTCF depleted cells. The domain score is also explained in right panel. The scores in non-synchronous and metaphase HFF-1 cells are also shown. The scores were preserved in CTCF depleted cells compared with metaphase HFF-1 cells in which TAD structures are disrupted completely. The domain score is also explained in right panel. (D) Scatter plots showing insulation scores at TAD boundaries. Each dot shows the average of insulation score for a bin that overlaps the TAD boundaries. Higher score denotes lower insulation. Increased insulation scores in varying degrees were observed in CTCF depleted cells, but overall the changes were mild. (E) Boundary strength is plotted for each boundary. Boundary strength is calculated by measuring the DI differential across each boundary as described in methods. The rows are ordered by boundary strength in the 0 hr sample for each clone, with the strongest boundaries at the top and weakest at the bottom. For each treatment duration (24hr, 48hr, 96hr), the column on the left shows boundary strength in that sample, and the column on the right shows the difference in boundary strength between that CTCF-depleted cells and the control cells. Density plots below each treatment duration show the distribution of values for the difference in boundary strength between treated and untreated cells. The percentage of boundaries that get weaker upon treatment is plotted to the left of the midline, and the percentage of boundaries that become stronger is plotted to the right of the midline. In each case, the vast majority of boundaries become weaker after auxin treatment. The bottom shows that the 2 clones have highly similar distributions of boundary strengths.

We next investigated whether the genome-wide loss of CTCF occupancy affects the higher order genome structure by principal component analysis of the Hi-C datasets, which reveals spatial segregation in compartments A and B harboring active and inactive chromatin, respectively (Lieberman-Aiden et al., 2009). Consistent with the above finding of no significant difference in the TAD structure (Figure 2A), the distributions of compartments A and B were remarkably similar before and after CTCF depletion in every time point (Figure S2C & S2D).

Previous studies have suggested that CTCF has critical roles in organizing genome (Ghirlando and Felsenfeld, 2016; Ong and Corces, 2014). Surprisingly, our results showed that the genome segmentations in TADs and compartment A/B were generally maintained in the absence of CTCF.

### Chromatin loops are weakened but persist in CTCF depleted cells

Previous studies have identified roughly 10,000 chromatin loops in various mammalian cell types, and a majority of them are anchored on CTCF binding sites preferentially organized in a head-to-head orientation (Guo et al., 2015; Rao et al., 2014; Sanborn et al., 2015). These CTCF-anchored chromatin loops have been shown to depend on CTCF binding sites, as deletion of the CTCF binding sites disrupts the loops (Narendra et al., 2016; Narendra et al., 2015; Sanborn et al., 2015). To examine the impact of CTCF depletion on chromatin loops we first used HICCUPS (Rao et al., 2014) to identify 4284 chromatin loops in untreated mESCs (FDR < 0.1%). As reported before, the majority of them (3199 peaks) were anchored at CTCF binding sites on both sides (Rao et al., 2014). We then performed Aggregate Peak Analysis (APA) (Rao et al., 2014) on the set of CTCF-anchored long-range loops (genomic distance > 100 kb) to compare the strength of these loops before and after CTCF depletion. The aggregated interaction frequency between loop anchors was noticeably reduced after auxin treatment but not completely abolished (Figure 3A). Surprisingly, the loop strength appeared to be regained after 96-hour treatment, when the number of CTCF peaks is only 1-2% of the initial number of peaks (Figure 3A). These results indicate that, while CTCF may be required for the robustness of chromatin loops, additional factors are likely involved in maintenance of the chromatin loops.

**Figure 3.**
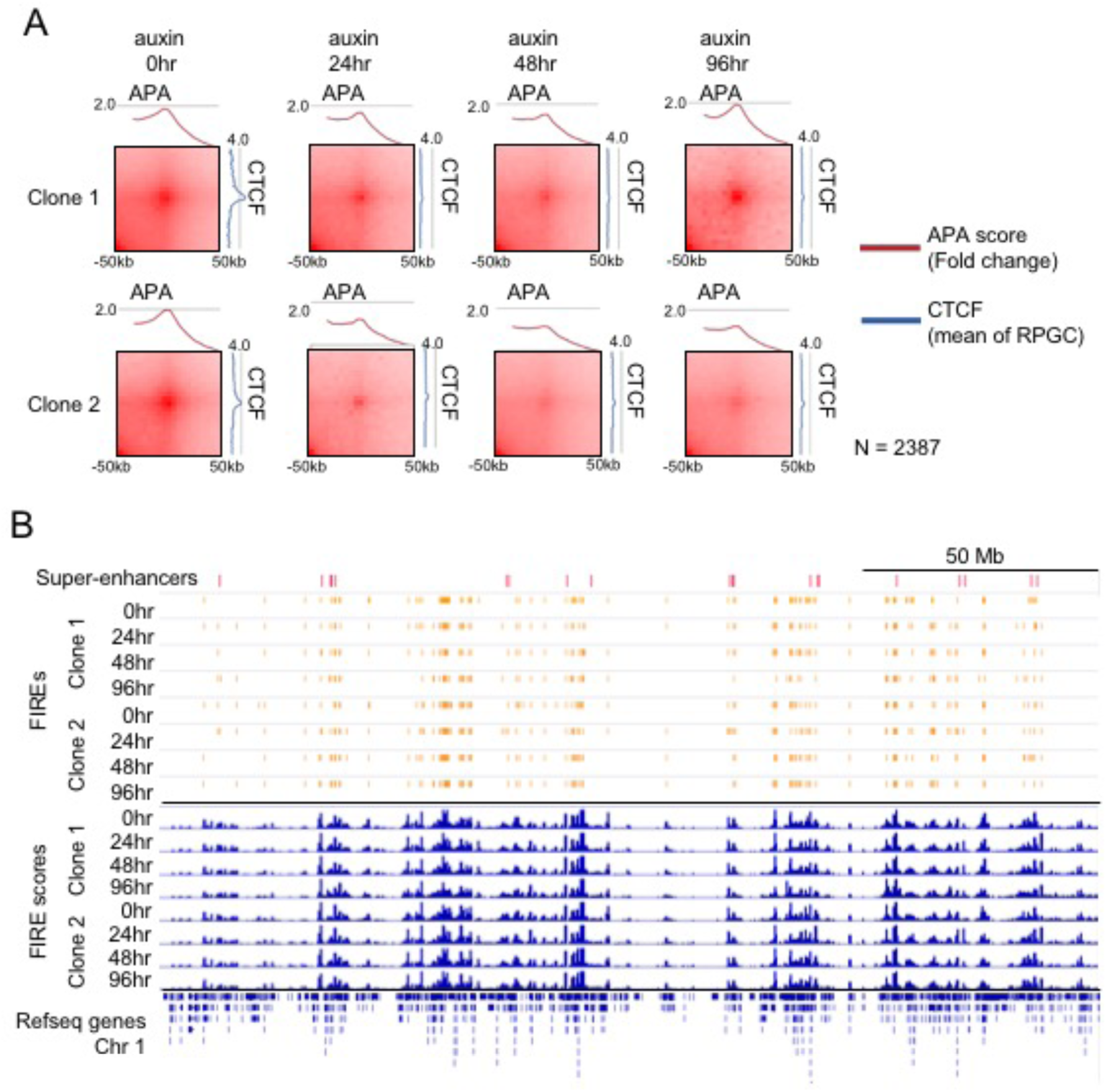
Chromatin loops are weakened but long-range chromatin interactions at enhancers in general are maintained after loss of CTCF. (**A**) CTCF-anchored chromatin loop that have over 100 kb range were identified in wild type mESCs (n=2387) and APA (aggregate peak analysis) was performed using the same loops in control and CTCD depleted cells in clone 1 and 2. Red line graph shows fold change of APA score. Blue line graph shows average of CTCF ChIP-seq signal. (**B**) Distributions of FIREs (orange bands) and FIRE scores (blue peaks) across entire chromosome 1 are stable to depletion of CTCF. Distributions of super enhancers and genes are shown at top and bottom respectively.

To confirm persistence of long-range chromatin interactions after CTCF depletion, we further investigated interactions around the *Sox2* gene using 4C-seq (Simonis et al., 2009). In mESC, *Sox2* is regulated by a super-enhancer located ~ 130 kb downstream of the gene (Li et al., 2014; Zhou et al., 2014). CTCF binding could be detected at the super-enhancer, and it was lost after the auxin treatment. The 4C-seq experiment showed that there is robust interaction between *Sox2* gene and the super-enhancer in untreated mESC, and the interaction persisted after days of auxin treatment (Figure S3A), supporting the idea that a CTCF-independent mechanism maintains the chromatin interactions between Sox2 and its downstream enhancer.

Recently, we reported frequently interacting regions (FIREs) as a feature of chromatin organization that are characterized by unusually high levels of local chromatin interactions and enrichment for active enhancers (Schmitt et al., 2016). We analyzed FIREs at each time point of auxin treatment (Figure 3B and Figure S3B). As for the case of TADs and compartments, the distribution and the number of FIREs were largely unchanged in CTCF depleted cells. The observations that FIREs are maintained after CTCF depletion are consistent with the results of the 4C-seq that revealed preservation of chromatin interactions between the *Sox2* gene and its downstream enhancer.

### Cohesin complex remains on chromatin temporarily after CTCF depletion

To explore the factors that may maintain TADs and chromatin loops in the absence of CTCF, we investigated the genomic distribution of Cohesin complex before and after CTCF depletion. Consistent with previous reports, ChIP-seq analysis of the Cohesin subunit Rad21 revealed a striking reduction of Cohesin binding genome-wide upon CTCF depletion (Figure 4A-B). Surprisingly, the rate at which Cohesin binding was lost was substantially slower than the loss of CTCF, and this difference was most apparent in Clone 1 (Figure 4B). The majority of Rad21 peaks in Clone 1 were still preserved after 24 hours of auxin treatment even though CTCF peaks had already been nearly completely lost. Rad21 occupancy then decreased progressively and reached the lowest number at the 96-hour time point. In Clone 2, the removal of Rad21 peaks was more rapid, with 95% of Rad21 occupancy lost after 24 hours of Auxin treatment (Figure 4C). The difference between the two clones might be caused by the different speeds of cell division (Figure 1C). These observations indicate that while the Cohesin complex requires CTCF for its genomic localization, as previously reported, it can remain associated with chromatin, albeit temporarily, after the removal CTCF protein. Since Cohesin occupancy is eventually lost from most TAD boundaries (Figure S4A), it is unlikely that Cohesin is responsible for maintaining TADs in the absence of CTCF (Figure S4B-C).

**Figure 4.**
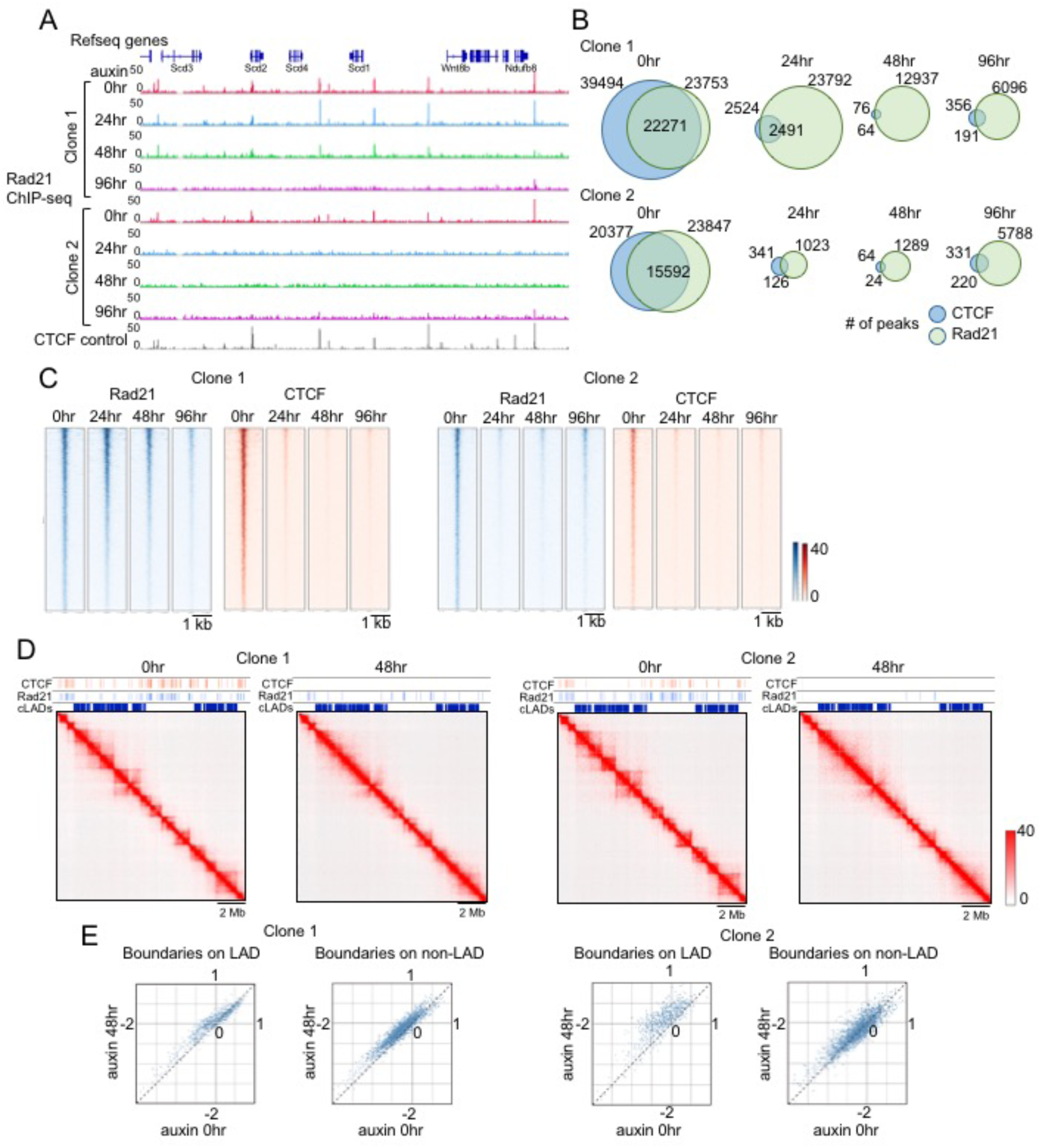
Cohesin complex is removed after CTCF depletion but at a slower pace than CTCF. (A) Snapshots of genome browser of two clones of Rad21 ChIP-seq experiments in control and CTCF depleted cells. In clone 1, the Rad21 peaks mostly remained after 24 hours of auxin treatment and then disappeared gradually after 48 and 96 hours of treatment. In clone 2, most of Rad21 peaks were removed by after 24 hours of auxin treatment. (B) Venn-diagrams showing the numbers of Rad21 ChIP-seq peaks in control and CTCF depleted cells along with the number of CTCF ChIP-seq peaks in clone 1 and 2. (C) Heatmaps comparing the Rad21 ChIP-seq signals centered at all regions of Rad21 peaks identified in the control cells and in CTCF-depleted cells at each time point (left, blue heat map) in clone 1 and 2. The CTCF occupancy at the same regions in each samples are also shown (right, red heat map). (D) Hi-C contact maps at 25 kb resolution from control cells and CTCF depleted cells (48 hours treatment of auxin) in clone 1 and 2. Distributions of CTCF and Rad21 ChIP-seq peaks and constitutive Lamina Associated Domains (cLADs) are aligned on their tops. (E) Scatter plots comparing insulation scores of bins at the TAD boundaries within cLADs and TAD boundaries outside cLADs.

### TAD structure within the Lamina-associated domains is sensitive to CTCF loss

A fraction of the genome is positioned near the nuclear membrane through association with the nuclear lamina and these genomic regions are referred to as lamina-associated domains (LADs) (Yanez-Cuna and van Steensel, 2017). LADs have been mapped in various mammalian cell types, and they generally correlate with inert transcription state and late replication timing (Gonzalez-Sandoval and Gasser, 2016; Yanez-Cuna and van Steensel, 2017). The constitutive LADs (cLADs) shared by all tested cell types are highly conserved in mouse and human and contain few genes (Meuleman et al., 2013; Peric-Hupkes and van Steensel, 2010). LADs and TADs often share the same boundaries (Dixon et al., 2012) and are also enriched for CTCF binding sites (Guelen et al., 2008; Meuleman et al., 2013). To investigate whether chromatin organization in cLADs are affected by loss of CTCF, we inspect the TAD structure within cLADs in the mouse ES cells. Interestingly, we found that TADs within these regions were frequently disrupted, in contrast to the TADs located outside LADs (Figure 4D). The insulation scores of the TAD boundaries in cLAD are more severely affected by loss of CTCF than TAD boundaries outside the LADs in both clones (Figure 4E). There was a small difference between the two cell clones with regard to the disruption of TADs by loss of CTCF. Clone 2 exhibited a more dramatic reduction of insulation score in TADs within cLADs than clone 1. This might be due to the faster growth rate of clone 2, which undergoes more cell divisions than clone 1 during 48 hours of auxin treatment (Figure 1C). Nevertheless, in both clones, the insulation scores at TAD boundaries outside the LAD regions are essentially unaffected by CTCF loss (Figure 4E). This result suggests that TAD structure in the LADs and outside LADs are differentially maintained, with the former likely more dependent on CTCF and the Cohesin complex than the latter.

### Gene expression profiles and chromatin state remain largely unaltered after CTCF depletion

In order to determine whether re-organization of TADs in the LADs leads to changes in gene expression, we performed RNA-seq with both clones before and 24hrs, 48hrs or 96 hrs after auxin treatment. Surprisingly, we identified fewer than 10 genes that were differentially expressed after 24 or 48 hours of auxin treatment (FDR < 0.1) (Figures 5A, S5A and Supplementary table 5). After 96 hours of auxin treatment, 253 genes were differentially expressed. We also performed ChIP-seq for three histone modifications marking active chromatin (H3K4me1, H3K4me3 and H3K27ac). In agreement with the lack of overall gene expression changes upon CTCF depletion, there was no significant change in chromatin modification patterns (Figure S5B). Consistent with a role for CTCF in gene regulation, the genes with CTCF binding at the transcription start sites (TSS) showed slightly more down-regulation in transcription than genes without CTCF binding sites at the TSS (Figures 5B and S5C). We also observed that genes in the LADs were affected by CTCF depletion to a higher degree than genes outside the LADs (Figure 5C). This finding, combined with the observation that TAD structure within LADs is more severely affected after loss of CTCF, points to a role for CTCF in regulating genome architecture and gene regulation within the specific context of LADs and not genome-wide, as has been widely assumed.

**Figure 5.**
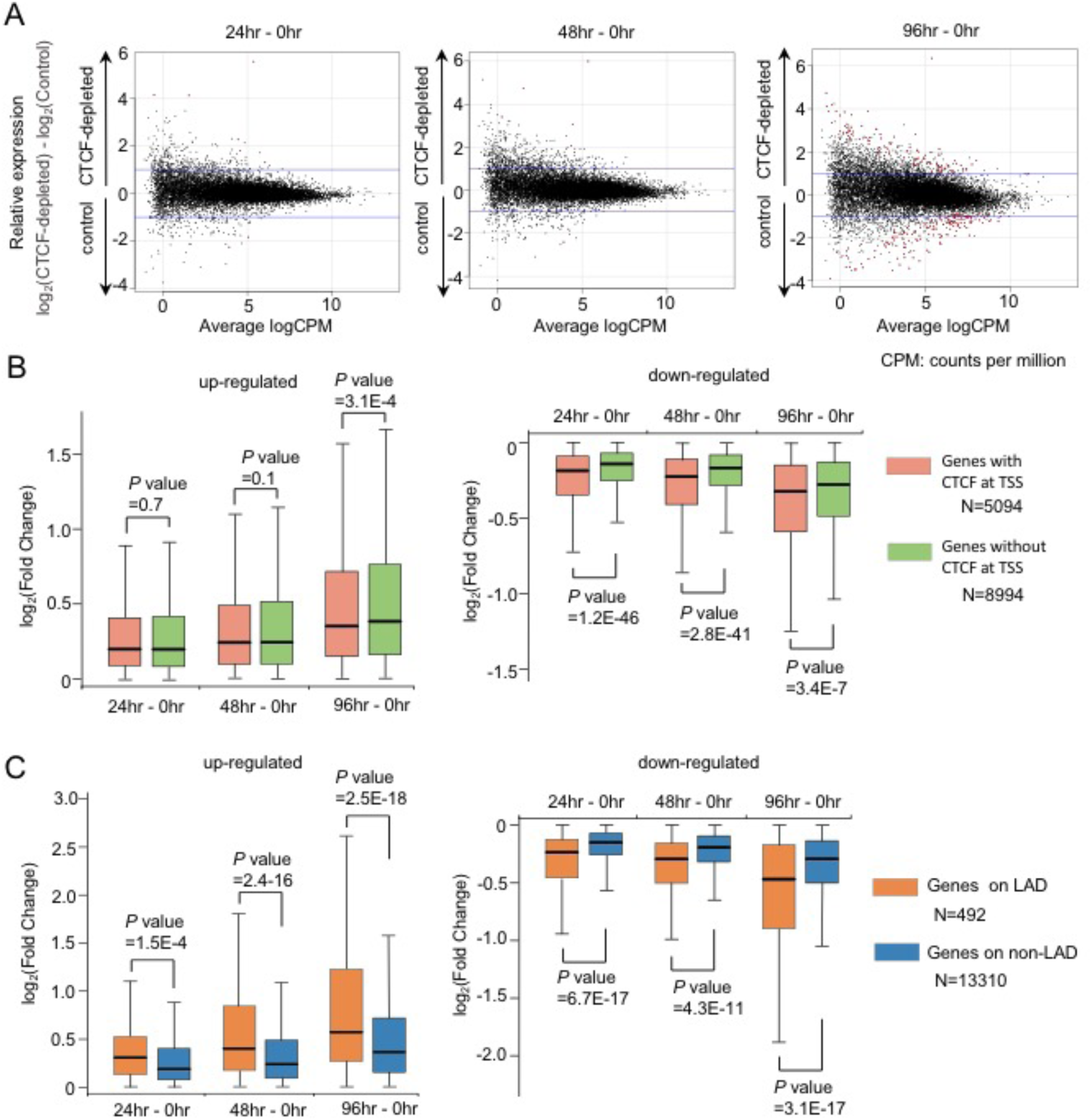
Transcription profiles are maintained in CTCF depleted cells. (A) RNA-seq MA plot of control versus CTCF-depleted cells treated with auxin for 24, 48, and 96 hours. Differentially expressed genes are showed as Red dots. Each panel is representative of two clones. (B) Box plots of gene expression changes in genes that have CTCF bindings at TSS prior to CTCF depletion (red) or not (green). The depletion of CTCF at TSS affected significantly in down-regulation of gene expression especially 24 and 48 hours after auxin treatment. Central bar, median; lower and upper box limits, 25th and 75th percentiles, respectively; whiskers, minimum and maximum value within the rage of (1st quartile-1.5*(3rd quartile-1st quartile)) to (3rd quartile+1.5*(3rd quartile-1st quartile)). (C) Box plots of gene expression changes in genes on cLADs (red) or on non-LAD (blue). Genes within LADs changed more significantly than genes outside of LADs (non-LAD) for both up- and down-regulated genes after CTCF depletion. Central bar, median; lower and upper box limits, 25th and 75th percentiles, respectively; whiskers, minimum and maximum value within the rage of (1st quartile-1.5*(3rd quartile-1st quartile)) to (3rd quartile+1.5*(3rd quartile- 1st quartile)).

### Discussion

The role of CTCF in chromatin organization in mammalian cells has been generally accepted, providing a framework for the understanding of long-range control of gene expression by *cis* regulatory elements such as enhancers and promoters. However, we demonstrate in this study that maintenance of higher order chromatin organization such as TADs and compartments does not strictly depend on CTCF. Using the auxin-inducible degron system, we acutely depleted CTCF protein in mouse embryonic stem cells, and monitored the impact of CTCF loss on genome architecture using Hi-C. We observed reduced chromatin looping strengths at CTCF-anchored chromatin loops but no overt changes in TAD structure and compartments, except for a small number of TADs located in constitutive LADs. We also failed to detect significant changes in chromatin states and transcription profiles. Our results suggest that the CTCF is not generally required for maintaining the TADs and compartmental organization of the genome, and additional factors must be involved in their demarcation.

Our results call for a revision of the current models of CTCF’s role in chromatin organization. One possible refinement is that there is a distinction between the mechanisms that establish chromatin loops at CTCF binding sites and the organization of chromatin into TADs and compartments. We observed that loss of CTCF in the mESC was indeed accompanied by significant reduction of chromatin looping anchored on CTCF binding sites, but the lack of substantial changes in TADs and compartments suggests that CTCF-mediated chromatin loops are not entirely responsible for the formation of TADs and compartments. This conclusion is consistent with previous findings that disruption of CTCF motifs led to loss of chromatin loops at certain CTCF binding sites but without changing the local chromatin domains (Sanborn et al., 2015). Also consistent with this conclusion is that CTCF binding sites are found more frequently within TADs than at TAD boundaries (Dixon et al., 2012). Furthermore, removal of the Cohesin complex in mammalian cells did not disrupt TADs (Seitan et al., 2013; Zuin et al., 2014). Thus, chromatin loops mediated by CTCF are insufficient to maintain TADs boundaries, and other mechanisms must be responsible. We note that in our experimental analysis, cell number at least doubled between 24h and 96h after auxin treatment (e.g. cell number increased ~4-fold for clone 2 between these time points). As TADs are disassembled in mitosis and reassembled during the transition to G1 (Dileep et al., 2015; Naumova et al., 2013), our results suggest that significant CTCF reduction also does not prevent TAD formation, although addressing this point will require additional future effort.

A second way to reconcile our observation and previous studies is that distinct mechanisms may be responsible for establishment of TADs in the late replicating LADs and early replicating non-LAD regions. As noted above, previous studies have shown that TADs are lost during mitosis and are re-established in early G1 (Dileep et al., 2015; Naumova et al., 2013). It is possible that different factors in different nuclear compartments might be involved in establishment or maintenance of TADs in the regions that replicate early and regions that replicate late during cell cycle. This could explain why the disruption of CTCF binding sites leads to the elimination of certain loops or TAD boundaries in LADs but not in non-LAD regions.

What mechanisms could be responsible for maintenance of TADs and chromatin compartments in the non-LAD regions? One potential factor is Topoisomerase II beta (Top2b), which has been found to interact with CTCF and Cohesin, and colocalizes with them at TAD boundaries (Uuskula-Reimand et al., 2016). Since Top2b can modulate the supercoiling of DNA, it is conceivable that TAD borders would have distinct topological property from inside the TAD, and such topological property is maintained after removal of CTCF. Another factor may be ZNF143, which like Top2b is colocalized with CTCF and Cohesin complex genome wide and at TAD borders (Bailey et al., 2015; Heidari et al., 2014). However, ZNF143 binds preferentially at promoters and may be primarily involved in mediating promoter-centered chromatin interactions. It is also possible that non-coding RNAs might be involved in establishing or maintaining TADs, as CTCF has been demonstrated to interact with many RNAs including the steroid receptor RNA activator SRA (Kung et al., 2015; Saldana-Meyer et al., 2014; Yao et al., 2010). Clearly, the role of CTCF in chromatin organization is not as clear-cut as previously proposed. Our study will help direct future efforts that promise to further elucidate the mechanisms of chromatin organization.

While this manuscript was under preparation, a similar work appeared in BioRxiv (Nora et al., doi:https://doi.org/10.1101/095802). With similar experimental approaches employed, the authors made similar observations with regard to the role of CTCF in chromatin loop formation and insulation scores at TAD boundaries. However, important distinctions exist between the two works with regard to TAD maintenance and transcriptional profiles after CTCF depletion. Future experiments are needed to address whether these differences were due to experimental data, data analysis methods or interpretation.

### Methods

Detailed methods and data analysis are described in the *supplemental method.*

## Acknowledgments

We would like to give special thanks to Samantha Kuan for operating the sequencing instruments. We would like to acknowledge the help of Ming Hu (NYU) for sharing software code. We would also like to give special thanks to Anthony Schmitt and Sora Chee for sharing protocols and giving numerous helpful advice, as well as the additional members of the Ren laboratory. This work was supported by the Ludwig Institute for Cancer Research (B.R.), NIH (1U54DK107977-01) (B.R.), NIH (1U54DK107965-01) (H.Z.) and a Postdoc fellowship from the TOYOBO Biotechnology Foundation (N.K.).

## Accession numbers

Sequencing data have been deposited in Gene Expression Omnibus (GEO) under accession number GSE94452, and can be accessed at https://www.ncbi.nlm.nih.gov/geo/query/acc.cgi?token=olctqyiktlyrhkf&acc=GSE94452.

## Author contributions

N.K., H.I. and B.R. conceived the project. H.I., X.X. and H.Z. engineered cell lines. N.K., T.L. and Z.Y. carried out library preparation. N.K., F.M. and A.D. performed cell cycle analysis. N.K., D.G., R.F. and B.L. performed data analysis. J.D. contributed to experimental design. N.K., H.I. and B.R. wrote the manuscript. All authors edited the manuscript.

## Supplemental Information

### 1. Supplemental Figures

**Figure S1.**
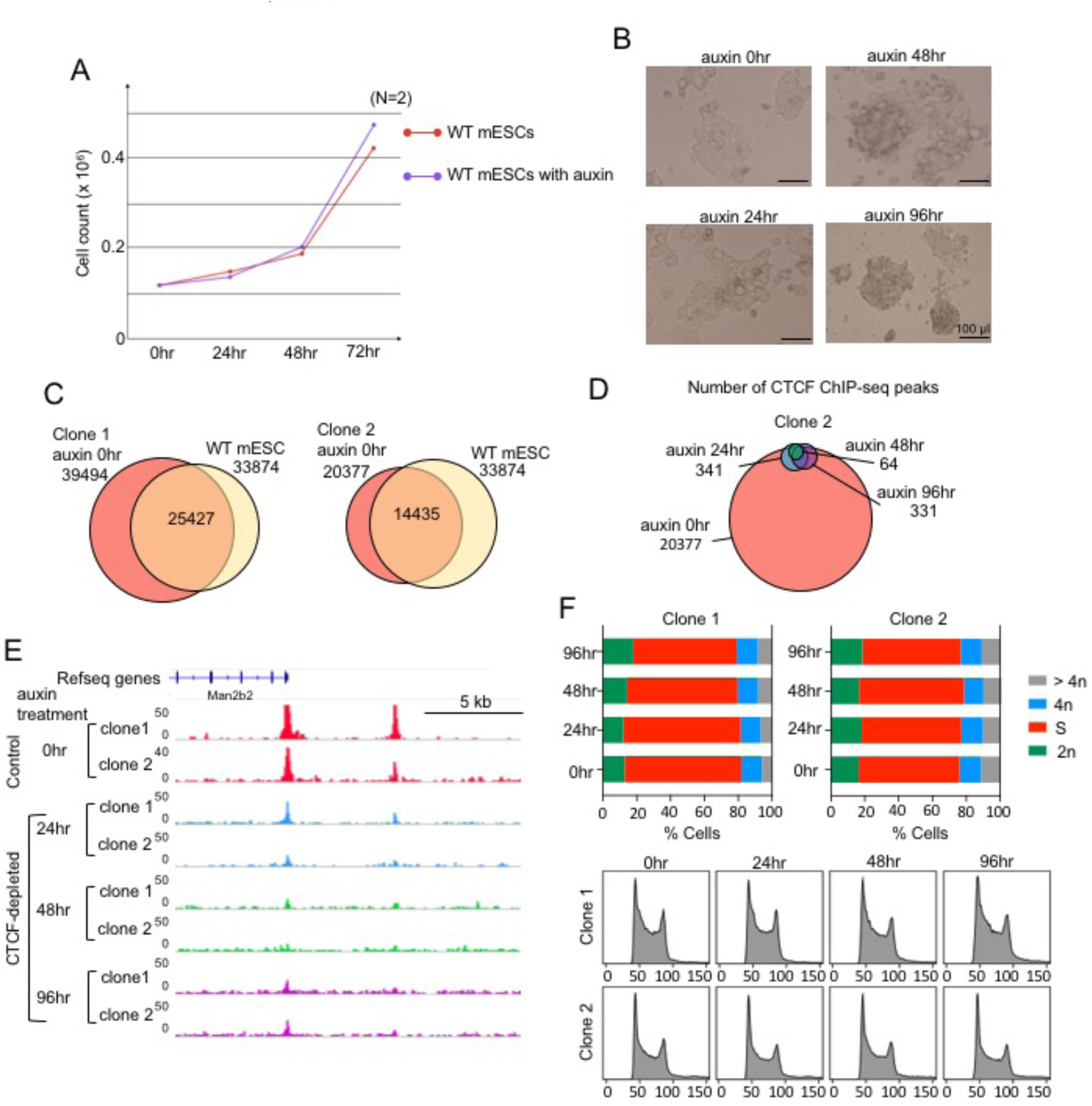
Characterization of the mESC after CTCF depletion. (**A**) Growth curves for mESC with or without treatment of Auxin. (**B**) Bright-field microscopy images of mESC colonies before and after auxin treatment. (**C**) Venn-diagram of the number of CTCF ChIP-seq peaks identified in the 2 clones of control cells and wild type mESCs and their overlapping. (**D**) Venn-diagram showing the numbers of CTCF ChIP-seq peaks identified in the control and CTCF depleted cells in clone 2. (**E**) An example of CTCF ChIP-seq peaks that were detected by peak calling in CTCF depleted cells. The enrichment level in these persistent peaks was severely reduced even though they were detected by peak calling. (**F**) Cell cycle analysis by flow cytometry using propidium iodide staining in control and 24, 48, and 96 hours CTCF-depleted cells indicates that CTCF depleted cells are not blocked in any stage of cell cycle.

**Figure S2.**
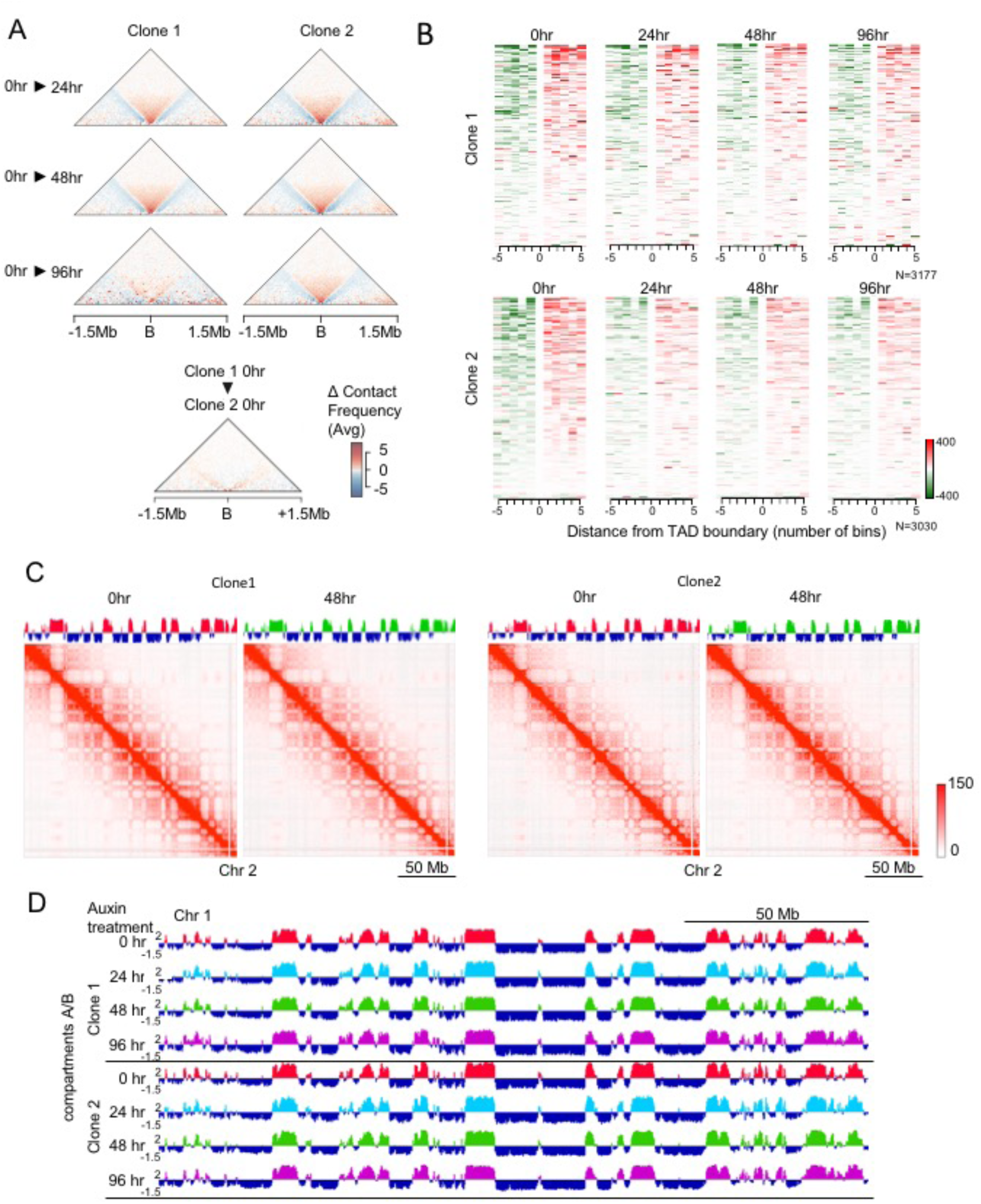
Chromatin organization is generally preserved after CTCF depletion. (**A**) Aggregate boundary analysis showing the average change in boundary strength between samples. Each triangle is a contact map showing the difference in the average contact profile at TAD boundaries between two time points in the auxin treatment regimen. The two time points being compared are indicated in the left margin. The top columns represent clones 1 and 2, respectively. The bottom column shows that there is little difference in the average boundary profile between clone 1 and clone 2 (without auxin treatment). B = Boundary. A consensus set of 4,364 boundaries were included in this analysis (see methods). (**B**) Directionality Index (DI) is plotted for 5 bins on either side of each TAD boundary. Rows are ordered by boundary strength in the 0 hr sample for each clone, with the strongest boundaries at the top and weakest at the bottom. For plotting the color scale was limited to -400:400. (**C**) Hi-C contact maps at 250 kb resolution across entire chromosome 2 from control cells and CTCF depleted cells (48 hours treatment of auxin) in clone 1 and 2. Distributions of compartment A/B are aligned on their tops. (**D)** Distributions of compartment A/B across entire chromosome 2 in clone 1 and 2.

**Figure S3.**
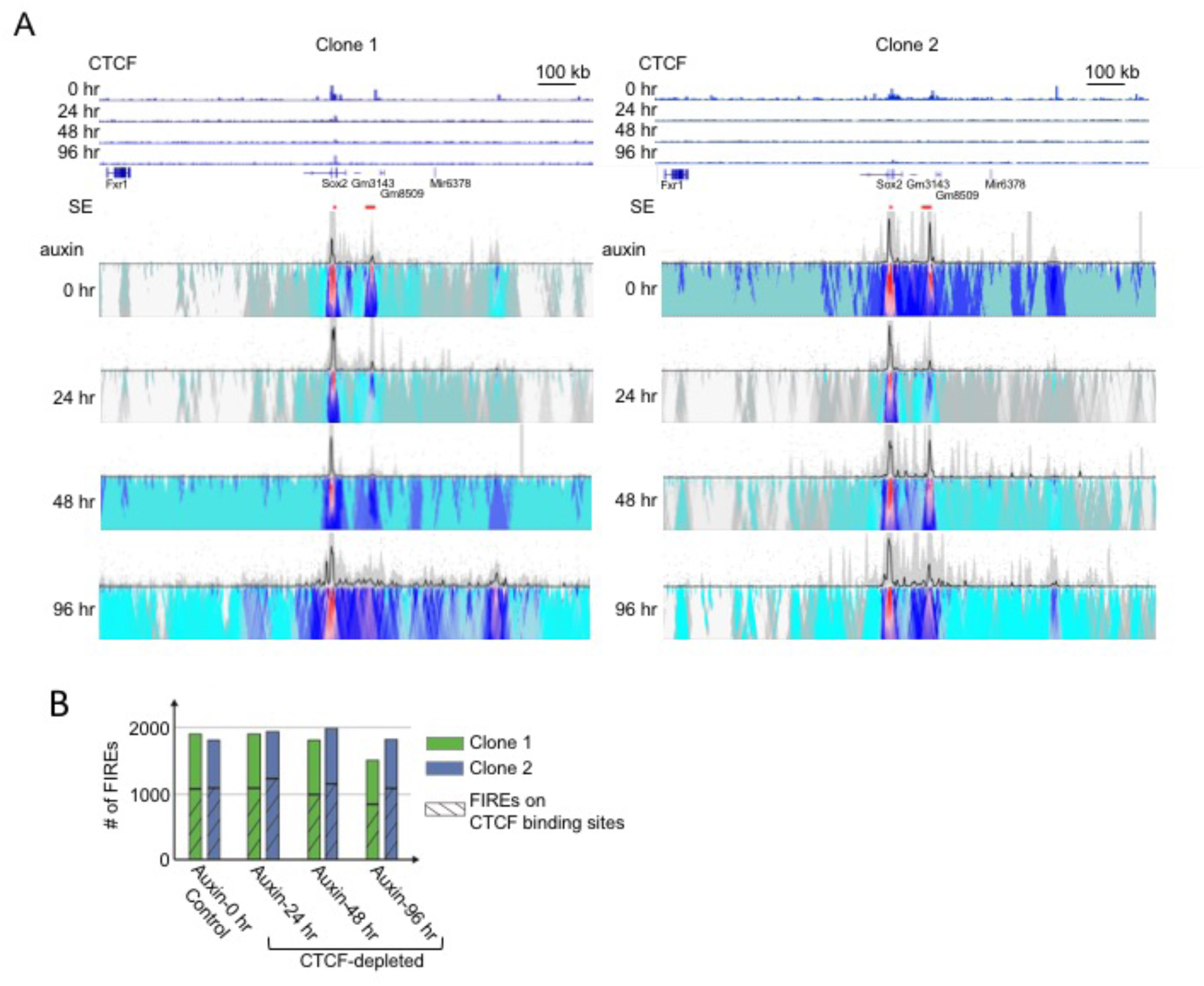
Chromatin organization at enhancers is not dramatically affected by CTCF loss. (**A**) 4C-seq showing chromatin interactions between *Sox2* promoter (used as bait) and its super enhancer (SE) after CTCF depletion in the both clones. CTCF ChIP-seq in each time point and distributions of SE (red band) are aligned on their top. (**B**) The numbers of FIREs in control and CTCF depleted cells in clone 1 and 2 are shown. The hatched bars indicate the number of FIREs that overlapped with CTCF ChIP-seq peaks.

**Figure S4.**
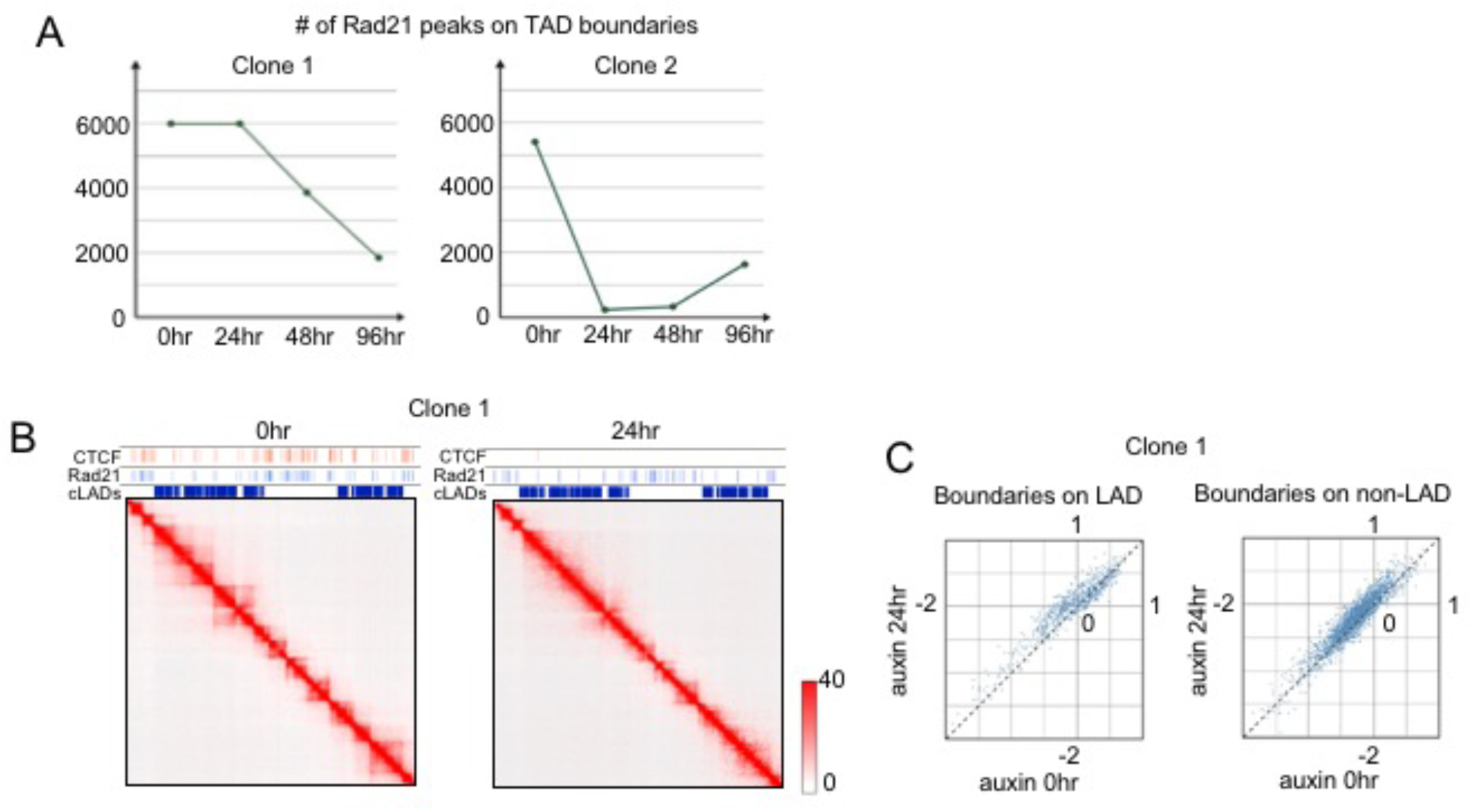
Cohesin binding and TAD structure are both affected by CTCF loss in the LADs. (**A**) The number of Rad21 peaks on TAD boundaries decreased after CTCF depletion in both clones. (**B**) Hi-C contact maps at 25 kb resolution from control cells and CTCF depleted cells (24 hours treatment of auxin) in clone 1. Distributions of CTCF and Rad21 ChIP-seq peaks and cLADs are aligned on their tops. In clone 1, 24 hours auxin treated cells still have many Rad21 peaks on cLADs, however the TAD structure is already disrupted on cLADs. (**C**) Scatter plots of insulation scores at TAD boundaries on cLADs and TAD boundaries on non-cLADs comparing control and 24 hours auxin treated cells. Almost all of the insulation scores on cLADs were increased even though Rad21 binding was largely maintained in those cells.

**Figure S5.**
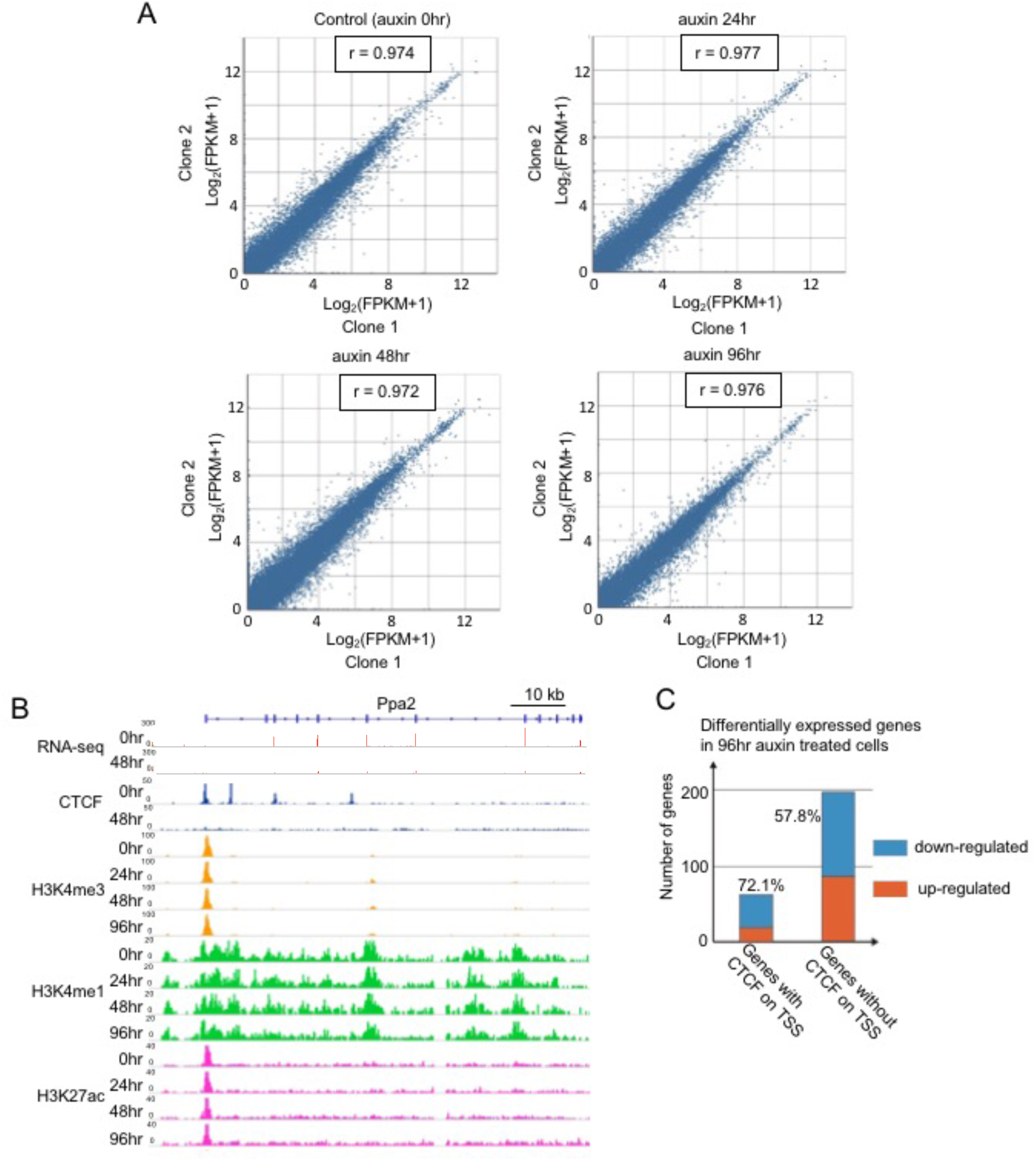
Transcription profiles and chromatin modifications are largely unchanged after CTCF loss. (**A**) Scatter plots showing high correlations between gene expression levels in clone 1 and those in 2 at every time point of auxin treatment. (**B**) Genome browser snapshot showing stable distribution of histone modifications (H3K4me3, H3K4me1 and H3K27ac) after CTCF depletion even around differentially expressed gene *Ppa2.* (**C**) Among the differentially expressed genes after 96 hour of auxin treatment (compared to control), the ratio of down-regulated genes are relatively high (72.1%) when they have CTCF binding on TSS prior to CTCF depletion.

### 2. SUPPLEMENTARY TABLES

**Table S1**. List of sequencing experiments and mapping summary, Related to Figure 1-5

**Table S2**. List of CTCF ChIP-seq peaks, Related to Figure 1

**Table S3**. List of TAD regions, Related to Figure 2

**Table S4**. List of Rad21 ChIP-seq peaks, Related to Figure 4

**Table S5**. List of differentially expressed genes, Related to Figure 5

### 3. Supplemental Methods and Materials

#### Construction of the plasmids

The CRISPR/Cas9 plasmid was assembled using the Multiplex CRISPR/Cas9 Assembly System (Sakuma et al., 2014) kit (a gift from Takashi Yamamoto, Addgene kit #1000000055). Oligonucleotides for three gRNA templates were synthesized, annealed and introduced into the corresponding intermediate vectors. The first gRNA matches the genome sequence 23 bp upstream of the stop codon of mouse CTCF. The oligonucleotides with sequences (5^’^- CACCGTGATCCTCAGCATGATGGAC-3’) and (5’-AAACGTCCATCATGCTGAGGATCAC-3^’^) were annealed. The other two gRNAs direct *in vivo* linearization of the donor vector: the first pair of oligonucleotides are (5’-CACCGCTGAGGATCATCTCAGGGGC -3^’^) and (5^’^- AAACGCCCCTGAGATGATCCTCAGC -3^’^); the second pair are (5^’^-CACCGATGCTGGGGCCTTGCTGGC-3’) and (5^’^-AAACGCCAGCAAGGCCCCAGCATC-3^’^). The three gRNA-expressing cassettes were incorporated into one single plasmid using Golden Gate assembly.

The donor vector was constructed using PCR and Gibson Assembly Cloning kit (New England Biolabs). The insert cassette includes sequences that codes for a 5GA linker, the auxin-induced degron (AID), a T2A peptide and the neomycin resistant marker, and is flanked by 24-bp homology arms to integrate into the CTCF locus. The left and right arms have sequences CCTGAGATGATCCTCAGCATGATG and GACCGGTGATGCTGGGGCCTTGCT, respectively. The AID coding sequence was amplified from pcDNA5-H2B-AID-EYFP(Holland et al., 2012) (a gift from Don Cleveland, Addgene plasmid #47329) and the T2A-neomycin was amplified from pAC95-pmax-dCas9VP160-2A-neo (Cheng et al., 2013) (a gift from Rudolf Jaenisch, Addgene plasmid #48227). The sequence for the 5GA linker was included in one of the primers. The original donor backbone was a gift from Dr. Ken-ichi T. Suzuki from Hiroshima University, Hiroshima, Japan.

The lentiviral vector for expressing TIR1 was constructed using PCR and Gibson Assembly Cloning kit (New England Biolabs). The backbone was modified from lentiCRISPR v2 (Sanjana et al., 2014) (a gift from Feng Zhang, Addgene plasmid #52961) and the TIR1-9myc fragment was amplified from pBabe TIR1-9myc (Holland et al., 2012) (a gift from Don Cleveland, Addgene plasmid #47328). The expressing cassette includes a puromycin resistant marker followed by sequences that code for P2A peptide and TIR1-9myc protein. The gene expression is driven by EFS promoter in the original lentiCRISPR v2.

#### Cell culture

The F1 Mus musculus castaneus × S129/SvJae mouse ES cells (F123 cells) (Gribnau et al., 2003) (a gift from Rudolf Jaenisch) were cultured in KnockOut Serum Replacement containing mouse ES cell media: DMEM 85%, 15% KnockOut Serum Replacement (Gibco), penicillin/streptomycin (Gibco), 1 × non-essential amino acids (Gibco), 1×GlutaMax (Gibco), 1000 U/ml LIF (Millipore), 0.4 mM β-mercaptoethanol. The cells were typically grown on 0.1% gelatin-coated plates with irradiated mouse embryonic fibroblasts (MEFs) (GlobalStem).

#### Transfection and establishment of CTCF-AID knock-in clones

The cells were passaged once on 0.1% gelatin-coated feeder-free plates before transfection. The cells were transfected using the Mouse ES Cell Nucleofector Kit (Lonza) and Amaxa Nucleofector (Lonza) with 10 μg of the CRISPR plasmid and 5 μg of the donor plasmid following the manufacturer’s instructions. After transfection, the cells were plated on drug-resistant MEFs (GlobalStem). Two days after transfection, drug selection was started by addition of 160 μg/ml G418 (Geneticin, Gibco) to the medium. Drug-resistant colonies were isolated and the clones with AID knock-in on both alleles were found by performing PCR of the genomic DNA using primers specific to sequences flanking the 3’ end of the CTCF coding sequence (AAATGTTAAAGTGGAGGCCTGTGAG and AAGATTTGGGCCGTTTAAACACAGC). The sequence at the CTCF-AID junction on both alleles were checked by sequencing of allele-specific PCR products, which were generated by using either a CTCF-129-specific (CTGACTTGGGCATCACTGCTG) or a CTCF-Cast-specific (GTTTTGTTTCTGTTGACTTAGGCATCACTGTTA) forward primer and a reverse primer in the AID coding sequence (GAGGTTTGGCTGGATCTTTAGGACA). The expression of CTCF-AID fusion protein was confirmed by observing the difference in the molecular weight compared to the control cells by Western blot with anti-CTCF antibody (Millipore, 07-729).

#### Lentivirus production and infection

We produced the lentivirus for expressing TIR1-9myc using Lenti-X Packaging Single Shots system (Clontech) and infected the CTCF-AID knock-in mouse ES cells following the manufacturer’s instructions. After infection, the cells were selected by culturing with 1 μg/ml puromycin. Drug-resistant colonies were isolated and expression of TIR1-9myc was confirmed by Western blot using anti-Myc antibody (Santa Cruz, sc-40). Clones expressing high level of TIR1-9myc were used for the subsequent experiments.

#### Preparation of CTCF-depleted cells

The CTCF-AID knock-in mouse ES cells expressing TIR1-9myc were passaged on 0.1% gelatin-coated plates without MEFs. We added 1 ul 500 mM auxin (Abcam, ab146403) per 1 ml medium to deplete CTCF, and changed medium with auxin every 24 hours. Cells were harvested 24, 48 or 96 hours after stating auxin treatment.

#### Antibodies

Primary antibodies used were rabbit anti-CTCF (Millipore, 07-729, for western blotting), rabbit anti-Histone H3 (abcam, western blotting), goat anti-CTCF (Santa Cruz, sc-15914, for microChIP-seq), rabbit anti-Rad21 (Santa Cruz, sc-98784, for microChIp-seq), rabbit anti-H3K4me1 (abcam, ab8895, for ChIP-seq), rabbit anti-H3K4me3 (Millipore, 04-745, for ChIP-seq) and rabbit anti-H3K27ac (Active Motif, 39133, for ChIP-seq).

#### Western blotting

mESCs were washed with PBS and scraped in cold PBS, and pelleted to be stored at -80°C. Two million cells were resuspended in 100 μL lysis buffer (20 mM Tris-HCl, 150 mM NaCl, 1 mM EDTA, 1mM EGTA, 1% Triton X-100, 1x complete protease inhibitor (Roche)), and sonicated for 10 minutes total ON time with pulses of 15 second ON and OFF, and 40% amplitude with QSONICA 800R (Qsonica). Protein concentration were measured using Pierce BCA Protein Assay Kit (Thermo Fisher). Laemmli Sample Buffer (Bio-Rad) with 355 mM 2-Mercaptoethanol were mixed with 15 μg of each sample and incubated for 5 minutes at 95°C. The samples were run on 4-15% Mini-PROTEAN^®^ TGX^™^ Precast Gels (Bio-Rad), and transferred onto nitrocellulose membranes at 100 V for 1 hour. The membranes were rinsed with 1x TBST and blocked with 5% dry milk at room temperature for 45 minutes. After washing with TBST, the membranes were incubated with diluted anti-CTCF antibody (Millipore, 07-729) (1:1000) in the blocking buffer overnight at 4°C. Membranes were washed 4 times 5minutes in 1x TBST at room temperature, incubated with secondary antibody, Goat Anti-Rabbit IgG (Bio-Rad) (1:2000), in blocking buffer at room temperature for 45 minutes. After washing 4 times with TBST, the substrates were detected using Pierce ECL Western Blotting Substrate (Thermo Fisher).

#### Cell cycle analysis

Cells were grown in 6-well plates. After dissociation with Accutase (Innovative Cell Technologies), 2-5 million cells were washed with PBS and re-suspended in 300 μl ice-cold PBS. Cells were fixed for a minimum of 24h at 4°C after drop-wise addition of 800 μl ice-cold ethanol. After fixation cells were pelleted and re-suspended in PBS containing 0.1% Triton X-100, 20 μg/mL Propidium iodide and 50 μg/ml RNase A. Cells were incubated for 30 min at 37°C before subjected to flow cytometry analysis.

#### MicroChIP-seq

MicroChIP-seq experiments for CTCF and Rad21 were performed as described in ENCODE experiments protocols (“Ren Lab ENCODE Chromatin Immunoprecipitation Protocol for MicroChIP” in https://www.encodeproject.org/documents/) with minor modifications. For fixation, trypsinized mES cells were counted and resuspended with adjusted volume (1 million cells per mL) of fresh media. Formaldehyde was added to 1% final, and samples were incubated for 10 minutes at room temperature. Then, 25 μl per mL of 2.5M Glycine was added, followed by a 5-minute incubation at room temperature and then a 15-minute incubation on ice. Aliquots were spun down at 3,500 x g for 15 minutes, washed with cold 1 x PBS, frozen on dry ice, and store at -80°C. We used 0.5 million cells for microChIP. Fixed cells were thawed on ice, and shearing of chromatin was performed using truChIP Chromatin Shearing Reagent Kit (Covaris) according to the manufacturer’s instructions. Covaris M220 was used for sonication with the following parameters: 10 minutes duration at 10.0% duty factor, 75.0 peak power, 200 cycles per burst at 5-9°C temperature range. The chromatin was diluted with 10 mM Tris-HCl pH 7.5, 140 mM NaCl, 1 mM EDTA, 0.5 mM EGTA, 1% Triton X-100, 10% SDS, 0.1% Sodium Deoxycholate, 1x complete protease inhibitor (Roche), 1 mM PMSF to adjust to 0.21% SDS concentration. The primary antibodies were prepared on magnetic beads as follows. We used 8 μL Magna ChIP Protein A+G Magnetic Beads (Millipore) for CTCF and 8 μL antirabbit IgG Dynabeads (Life Technologies) for Rad21. We washed the beads with cold RIPA buffer 1 (10 mM Tris-HCl pH 7.5, 140 mM NaCl, 1 mM EDTA, 0.5 mM EGTA, 1% Triton X-100, 0.1% SDS, 0.1% Sodium Deoxycholate) for 2 times. After washing, 5 μL anti-CTCF antibody (Santa Cruz, sc-15914) or 5 μL anti-Rad21 (Santa Cruz, sc-98784) with 95 μL RIPA buffer 1 were added to the beads and incubated on a rotating platform at 4°C for 6 hours. After incubation, beads were washed once with 100 μL cold RIPA buffer 1, and mixed with chromatin followed by overnight incubation on a rotating platform at 4°C. Beads were collected on a magnetic rack and washed 4 times with 10 mM Tris-HCl pH 7.5, 300 mM NacCl, 1 mM EDTA, 0.5 mM EGTA, 1% Triton X-100, 0.2% SDS, 0.1% Sodium Deoxycholate and washed once with 100 μL cold 1x TE. After removing the TE, 150 μL elution buffer 1 (20 mM Tris-HCl pH 7.5, 50 mM NacCl, 5 mM EDTA, 1% SDS) was added. For input samples, elution buffer 1 was also added to the chromatin set aside after the sonication to adjust the total volume to 300 μL. The input samples were processed in parallel with the ChIP samples from here on. RNase A (final conc. = 0.2 mg/mL) was added to each sample and incubated at 37°C for 1 hour, with shaking at 1200rpm on a thermomixer. Then, proteinase K (final conc. = 0.13 mg/mL) was added and the samples were incubated at 68°C for 4 hours, shaking at 1200rpm on a thermomixer. The beads were removed using a magnetic rack. The samples were extracted with Phenol: Chloroform: Isoamyl Alcohol (25:24:1) using Phase Lock tubes (5 Prime). Then the samples were precipitated with ethanol and resuspended in 25 μL 10 mM Tris. To prepare Illumina sequencing libraries, ThruPLEX DNA-seq 12s kit (Rubicon Genomics) was used according to the manufacturer’s instructions. We used 0.5-1.0 ng IP materials and 50 ng input DNA for library preparation, and 11-12 and 5 cycles of PCR were performed respectively. After purification by 1x AMpure Beads (Beckman Coulter), library quality and quantity were estimated with TapeStation (Agilent Technologies) and Qubit (Thermo Fisher Scientific) assays. Libraries were sequenced on HiSeq2500 or HiSeq4000 single end for 50 bp.

#### ChIP-seq

ChIP-seq experiments was performed as described in ENCODE experiments protocols (“Ren Lab ENCODE Chromatin Immunoprecipitation Protocol” in https://www.encodeproject.org/documents/) with minor modifications. Cells were crosslinked with 1% formaldehyde for 10 minutes and quenched with 125mM glycine. We used 1.0 million cells for each ChIP sample. Shearing of chromatin was performed using truChIP Chromatin Shearing Reagent Kit (Covaris) according to the manufacturer’s instructions. Covaris M220 was used for sonication with following parameters: 10 minutes duration at 10.0% duty factor, 75.0 peak power, 200 cycles per burst at 5-9°C temperature range. The concentration of fragmented DNA was measured using NanoDrop and diluted to 0.2 μg/μl with 1x TE. For immunoprecipitation, we used 11 μL anti-rabbit IgG Dynabeads (Life Technologies) and wash them with cold BSA/PBS (0.5 mg / mL bovine serum albumin in 1x phosphate buffered saline) for 3 times. After washing, 3 μL anti-H3K4me1 (abcam, ab8895), anti-H3K4me3 (Millipore, 04-745) or anti-H3K27ac (Active Motif, 39133) with 147 μL cold BSA/PBS were added to the beads and incubated on a rotating platform at 4°C for 2 hours. After incubation, beads were washed with150 μL cold BSA/PBS for 3 times, and mixed with 100 μL Binding Buffer (1% Triton X-100, 0.1% Sodium Deoxycholate, 1x complete protease inhibitor (Roche)) plus 100 μL 0.2 μg/μl chromatin followed by overnight incubation on a rotating platform at 4°C. Beads were collected on a magnetic rack and washed 5 times with 50 mM Hepes pH 8.0, 1% NP-40, 1 mM EDTA, 0.70% Sodium Deoxycholate, 0.5 M LiCl, 1x complete protease inhibitor (Roche) and washed once with 150 μL cold 1x TE. After removing the TE, 150 μL ChIP elution buffer (10 mM Tris-HCl pH 8.0, 1 mM EDTA, 1 % SDS) was added, and samples were incubated at 65°C for 20 minutes at 1300 rpm on a thermomixer. The beads were removed using a magnetic rack and the samples were further incubated at 65°C overnight to reverse crosslinks. For input samples, 20 μL of chromatin was added to 130 μL ChIP elution buffer and incubated at 65°C overnight with the other samples. The input samples were processed in parallel with the ChIP samples from here on. RNase A (final conc. = 0.2 mg/mL) was added to each sample and incubated at 37°C for 1 hour, and Proteinase K (final conc. = 0.4 mg/mL) was added and incubated at 55°C for 1 hour. The samples were extracted with phenol: chloroform: isoamyl alcohol (25:24:1) using Phase Lock tube (5 Prime). Then the samples were precipitated with ethanol and resuspended in 50 μL 10 mM Tris pH 8.0. We used 3-5 ng of starting IP materials for preparing Illumina sequencing libraries. The End-It DNA End-Repair Kit (Epicentre) was used to repair DNA fragments to blunt ends, and Qiagen MinElute PCR Purification kit (Qiagen, Cat#28006) was used to purify the samples. A-tailing 3^’^ end was performed using Klenow Fragment (3’ → 5’ exo-) (New England Biolabs), and then TruSeq Adapters were ligated using Quick T4 DNA Ligase (New England Biolabs). The samples were run on 2% agarose gel, gel slices corresponding to 300-500bp DNA were cut, and the DNA was purified using Qiagen MinElute Gel Extraction kit (Qiagen). After deciding optimal PCR cycle number using KAPA DNA Quantification kit (Kapa Biosystems), 8-10 cycles of PCR amplification was performed using KAPA Library Amplification kit (Kapa Biosystems). After purification by 1x AMpure Beads (Beckman Coulter), Library quality and quantity were estimated with TapeStation (Agilent Technologies) and Qubit (Thermo Fisher Scientific) assays. Libraries were sequenced on HiSeq4000 single end for 50 bp.

#### RNA-seq

Total RNA was extracted from 1-2 million cells using the AllPrep Mini kit (QIAGEN) according to the manufacturer’s instructions and 1 μg of total RNA was used to prepare each RNA-seq library. The libraries were prepared using TruSeq Stranded mRNA Library Prep Kit (Illumina). Library quality and quantity were estimated with TapeStation (Agilent Technologies) and Qubit (Thermo Fisher Scientific) assays. Libraries were sequenced on HiSeq4000 using 50 bp paired end.

#### Hi-C

*In situ* Hi-C experiments were performed as previously described using the MboI restriction enzyme (Rao et al., 2014). For fixation, trypsinized mES cells were counted and resuspended with adjusted volume (1 million cells per mL) of fresh media. Formaldehyde was added to 1 % final, and samples were incubated for 10 minutes at room temperature. Then, 25 μl per mL of 2.5M Glycine was added followed by a 5-minute incubation at room temperature and then a 15-minute incubation on ice. Aliquots were spun down at 3,500 x g for 15 minutes, washed with cold 1 x PBS, frozen on dry ice, and stored at -80°C.

The crosslinked pellets were thawed on ice, and were incubated with 200ul of lysis buffer (10 mM Tris-HCl pH 8.0, 10 mM NaCl, 0.2% Igepal CA630, 33 μL Protease Inhibitor (Sigma, P8340)) on ice for 15 min, washed with 300 μL cold lysis buffer, and then incubated in 50uL of 0.5% SDS for 10min at 62°C. After heating, 170 μL of 1.47% Triton X-100 was added and incubated for 15min at 37°C. To digest chromatin 100U MboI and 25uL of 10X NEBuffer2 were added followed by overnight incubation at 37°C with agitation at 700rpm on a thermomixer. After incubation, MboI was inactivated by heating at 62°C for 20 minutes. Digestion efficiency was confirmed by performing agarose gel electrophoresis of the samples.

The digested ends were filled and labeled with biotin by adding 37.5uL of 0.4mM biotin-14-dATP (Life Tech), 1.5 μL of 10mM dCTP, 10mM dTTP, 10mM dGTP, and 8uL of 5U/ul Klenow (New England Biolabs) and incubating at 23°C for 60 minutes with shaking at 500 rpm on a thermomixer. Then the samples were mixed with 1x T4 DNA ligase buffer (New England Biolabs), 0.83% Trition X-100, 0.1 mg/mL BSA, 2000U T4 DNA Ligase (New England Biolabs, M0202), and incubated for at 23°C for 4 hours with shaking at 300rpm on a thermomixer to ligate the ends. After the ligation reaction, samples were spun and pellets were resuspended in 550uL 10 mM Tris-HCl, pH 8.0. To digest the proteins and to reverse the crosslinks, 50 μL of 20mg/mL Proteinase K (New England Biolabs) and 57 μL of 10% SDS were mixed with the samples, and incubated at 55°C for 30 minutes, and then 67 μL of 5M NaCl were added followed by over night incubation at 68°C.

After cooling the samples for 10 minutes at room temperature, 0.8X Ampure (Beckman-Coulter) purification was performed and samples were eluted in 100 μL 10 mM Tris-HCl, pH 8.0. Next, the samples were sonicated to mean fragment length of 400 bp using Covaris M220 with the following parameters: 70 seconds duration at 10.0% duty factor, 50.0 peak power, 200 cycles per burst. To collect 200-600 bp size of fragmented DNA, two rounds of Ampure (Beckman-Coulter) beads purification was performed and the samples were eluted in 300 μL 10 mM Tris-HCl, pH 8.0.

The DNA labeled with biotin was purified using Dynabeads My One T1 Streptavidin beads (Invitrogen). 100 μL of 10 mg/mL Dynabeads My One T1 Streptavidin beads was washed with 400 μL of 1x Tween Wash Buffer (5 mM Tris-HCl pH 7.5, 0.5 mM EDTA, 1 M NaCl, 0.05% Tween-20), and resuspended in 300 μL of 2x Binding Buffer (10 mM Tris-HCl pH 7.5, 1 mM EDTA, 2 M NaCl). The beads were transferred to the sample tube, incubated for 15 minutes at room temperature, and the supernatant was removed from the beads using a magnetic rack. Then the beads were washed twice by adding 600 μL of 1x Tween Wash Buffer, heating on a thermomixer for 2 minutes at 55°C with mixing, and removing the supernatant using a magnetic rack. Then the beads were equilibrated once in 100 uL 1x NEB T4 DNA ligase buffer (New England Biolabs) followed by removal of the supernatant using a magnetic rack. To repair the fragmented ends and remove biotin from unligated ends, the beads were resuspended in 100uL of the following: 88 μL 1X NEB T4 DNA ligase buffer (New England Biolabs, B0202), 2 μL of 25mM dNTP mix, 5 μL of 10 U/μL T4 PNK (New England Biolabs), 4 μL of 3 U/μL NEB T4 DNA Polymerase (New England Biolabs), 1 μL of 5U/μL Klenow (New England Biolabs). The beads were incubated for 30 minutes at room temperature, followed by removal of the supernatant using a magnetic rack. The beads were washed twice by adding 600 μL of 1x Tween Wash Buffer, heating on a thermomixer for 2 minutes at 55°C with mixing, and removing the supernatant using a magnetic rack. To add dA-tail, the beads were resuspended in 100uL of the following: 90 μL of 1X NEB Buffer2, 5 μL of 10mM dATP, and 5 μL of 5U/ul Klenow (exo-) (New England Biolabs). The beads were incubated for 30 minutes at 37°C, followed by removal of the supernatant using a magnetic rack. The beads were washed twice by adding 600 μL of 1x Tween Wash Buffer, heating on a thermomixer for 2 minutes at 55°C with mixing, and removing the supernatant using a magnetic rack. Following the washes, the beads were equilibrated once in 100 μL 1x NEB Quick Ligation Reaction Buffer (New England Biolabs) and the supernatants were removed using a magnetic rack. Then the beads were resuspended again in 50 μL 1x NEB Quick Ligation Reaction Buffer. To ligate adapters, 2 μL of NEB DNA Quick Ligase (New England Biolabs) and 3 μL of Illumina Indexed adapter were added to the beads, and incubated for 15 minutes at room temperature. The supernatant was removed using a magnetic rack and the beads were washed twice by adding 600 μL of 1x Tween Wash Buffer, heating on a thermomixer for 2 minutes at 55°C with mixing, and removing the supernatant using a magnetic rack. Then the beads were resuspended once in 100 μL 10 mM Tris-HCl, pH 8.0, followed by removal of the supernatant and resuspension again in 50 μL 10 mM Tris-HCl, pH 8.0. After deciding an optimal PCR cycle number using KAPA DNA Quantification kit (Kapa Biosystems), 8-9 cycles of PCR amplification was performed with the following: 10 μL Fusion HF Buffer (New England Biolabs), 3.125 μL 10uM TruSeq Primer 1, 3.125 μL 10uM TruSeq Primer 2, 1 μL 10mM dNTPs, 0.5 μL Fusion HotStartII, 20.75 μL ddH20, 11.5 μL Bead-bound HiC library. Then, PCR products underwent final purification using AMPure beads (Beckman-Coulter), and were eluted in 30 μL 10 mM Tris-HCl, pH 8.0. Libraries were sequenced on Illumina HiSeq 4000.

#### 4C-seq experiments and data analysis

4C-seq experiments and data analysis were performed following the the procedures described previously (van de Werken et al., 2012). 4C-seq was performed on lysed 1% crosslinked pellet. Lysed cells were digested with Csp6I for the first restriction enzyme digestion, ligated with T4 DNA ligase, digested with NlaIII for the second restriction digestion, and ligated again. DNA was then purified using isopropanol precipitation and cleaned-up using AMPure beads (Beckman-Coulter). PCR reactions were performed using 1 μg of 4C template for 30 cycles in the bait region. PCR products were purified using Roche’s High Pure PCR Product Purification kit. Second round of 10 PCR cycle was performed using 1 μg of purified PCR products to anneal Illumina’s TruSeq adaptors necessary for sequencing. PCR products underwent final purification using AMPure beads (Beckman-Coulter). Libraries were combined and spiked into other non-4C seq libraries and sequenced on Illumina HiSeq 4000.

#### ChIP-seq data analysis

Each fastq file was mapped to mouse genome (mm10) with Bowtie (Langmead et al., 2009) allowing 1 mismatch using -m 1 option. The bigWig files were created using deepTools (Ramírez et al., 2016) with following parameters: --ignoreDuplicates --normalizeTo1x 2150570000 -- ignoreForNormalization chrX. The deepTools was also used for generating heat maps in figure 1 and 4. PCR duplicate reads were removed using Picard MarkDuplicates before peak calling and peaks were called with input control using MACS (Zhang et al., 2008) with default parameters.

#### RNA-seq data analysis

RNA-seq reads (paired end, 100 bases) were aligned against the mouse mm10 genome assembly using TopHat (Trapnell et al., 2009). The mapped reads were counted using HTSeq (Anders et al., 2015) and the output files from two replicates were subsequently analyzed by edgeR to estimate the transcript abundance and to detect the differentially expressed genes (Robinson et al., 2010). Differentially expressed genes were called by an FDR <10%. To analyze correlation between two replicates, the mapped reads from each replicate were subsequently analyzed by Cufflinks to estimate the transcript abundance (Trapnell et al., 2012), and the expression level indicated as FPKM was used for correlation analysis.

#### Hi-C Data analysis

Hi-C reads (paired end, 50 or 100 bases) were aligned against the mm10 genome using BWA-mem. To remove the short-range Hi-C artifacts - unligated and self-ligated Hi-C molecules - we removed reads whose paired mapping distance was less than 15 kb. PCR duplicate reads were removed using Picard MarkDuplicates. Raw contact matrices were constructed using in-house scripts with 40 kb resolution, and then normalized using HiCNorm (Hu et al., 2012). We used juicebox pre (Durand et al., 2016) to create hic file with -q 30 -f options. To visualize Hi-C data, we used Juicebox (Durand et al., 2016) and 3D Genome Browser (http://www.3dgenome.org). Topological domain boundaries were identified at 40-kb resolution based on the directionality index (DI) score and a Hidden Markov Model as previously described (Dixon et al., 2012). The insulation score analysis was performed as previously described (Crane et al., 2015) and insulation scores on TAD boundaries were calculated by taking average value of scores that overlapped with TAD boundaries. Compartment A/B analysis was performed at 100 kb resolution using the HOMER tool with default parameters (Heinz et al., 2010), and frequently interacting regions (FIREs) analysis were performed as previously described (Schmitt et al., 2016).

To assess global changes in TAD boundary strength between samples, we performed a comparison of each samples’ aggregated boundary contact profile. First, to generate a consensus set of TAD boundaries we performed a simple merge between boundaries from clone 1 prior to auxin treatment (Clone 1, 0 hr) and boundaries from clone 2 prior to auxin treatment (Clone 2, 0 hr). Two filtering steps were used to generate the final set of consensus boundaries: 1) We discarded boundaries there were within 3.04 Mb of a chromosome start or end, because we would not be able to extract a submatrix of the correct size for the aggregate analysis; 2) We discarded boundaries > 200kb, because these often represent regions of disorganized chromatin between TADs, rather than true TAD boundaries. Next, we extracted a Hi-C sub-matrix for each boundary in each sample. Each sub-matrix consists of a window of 3.04 Mb centered on the midpoint of the boundary in question. These boundary sub-matrices were then averaged to generate one 3.04 Mb matrix representing the average boundary contact profile in a given sample. To facilitate comparison between samples, average boundary contact profiles were then normalized across samples using standard quantile normalization. We then made pairwise comparisons between samples by subtracting the average boundary contact profile of sample 1 from the average boundary contact profile of sample 2. The list of consensus TAD boundaries used here is same as that described for the aggregate boundary analysis above. To quantify the strength of each boundary, we calculated the difference in DI across each boundary using a simple formula: Boundary strength = (mean of DI values in 3 bins to the right of boundary) - (mean of DI values in 3 bins to the left of boundary).

To identify chromatin loops in our engineered untreated mESCs, we used HiCCUPS (Durand et al., 2016) with 5 kb and 10 kb resolutions and option “-r 5000,10000 -f 0.001,0.001 -p 4,2 -i 7,5 -d 20000,20000”, and then we chose CTCF associated loops among them that were overlapped with CTCF ChIP-seq peaks in control cells and other peaks in mESCs from public database on both sides of looping interaction. The aggregate analysis of CTCF associated loops were performed using APA (Durand et al., 2016) with default parameters except for “-n 5” option.

#### Data from other sources

CTCF ChIP-seq in wild type mES cells were downloaded from GSE39502 (Plasschaert et al., 2014) and GSE36027 (ENCODE). The data of Constitutive lamina associated domains was downloaded from GSE17051 (Peric-Hupkes et al., 2010), and the data of super-enhancer was taken from dbSUPER database (Khan and Zhang, 2016).

